# Human adipose-derived mesenchymal stromal cells improved wound healing in enterocutaneous fistulizing disease mouse model

**DOI:** 10.1101/2024.11.07.622127

**Authors:** Noga Marsiano, Yeshurun Levi, Peleg Schneider, Iftach Schouten, Einat Nisim-Eliraz, Simcha Yagel, Kok-Fai Kong, Eleuterio Lombardo, Nahum Y. Shpigel

## Abstract

A significant complication of Crohn’s disease is the formation of perianal fistulas. While local application of human adipose tissue-derived mesenchymal stromal cells (AT-MSC INN Darvadstrocel, Takeda) is an approved therapy for this condition, challenges in studying human clinical tissue and the absence of a robust experimental model have hindered deeper understanding of the preclinical efficacy and mechanisms behind the therapy. We have developed a model that closely mimics the clinical scenario, using human gut tissue transplanted subcutaneously into SCID mice. In this model, enterocutaneous fistulas are reliably induced by combining imiquimod-induced psoriatic dermatitis above the transplant with systemic lipopolysaccharide (LPS), a major component of the bacterial cell wall. Similar to clinical observations, local application of AT-MSC significantly improved fistula wound healing and re-epithelialization. This model system enabled comprehensive tissue harvesting for histopathological analysis, spatial transcriptomics, and protein profiling.

Our findings demonstrate that AT-MSC applied around the fistula wound survive for up to three weeks, migrating into the fistula tract and further into the inflamed human gut tissue. Within the fistula tract, AT-MSCs acquired immune-active properties, with increased expression of SOD2 and CCL2, and were associated with a substantial population of M2 macrophages. In contrast, AT-MSC located in healthy tissue near the fistula remained stationary, adopting a fibroblast-like phenotype within a collagen-rich extracellular matrix. These preclinical results support the safety and efficacy of AT-MSC for treating fistulizing gut disease. Furthermore, our data suggest that the inflamed fistula tract acts as a scaffold promoting AT-MSC activity and migration. Enhancing SOD2 and CCL2 expression through pre-activation or genetic modification may further improve the therapeutic potential of these cells.

## INTRODUCTION

Inflammatory Bowel Disease (IBD) is an incurable condition that is increasing in incidence, affecting nearly 3 million young adults in the United States (US) alone (Lewis et al., 2023). Crohn’s Disease (CD) accounts for many of these patients, causing significant morbidity. The aetiology of CD is unknown but is likely driven by aberrant adaptive immune responses to microbial antigens in the context of a permissive genetic background (Guan, 2019; Petagna et al., 2020; Yuan et al., 2024). In essence, a loss of tolerance to non-self antigens causes pathogenic effector T cells to induce recurrent bouts of inflammation throughout the entire gastrointestinal tract, from mouth to anus. Current treatments work by blocking specific cytokines or T cell signalling/trafficking pathways utilized by pathogenic T cells and act to block these molecular interactions rather than cure disease. Perianal fistulizing CD (fCD), is a particular unmet medical need (Anandabaskaran et al., 2023; Kotze et al., 2018; Lightner et al., 2020; Tozer et al., 2018; White et al., 2024). It is more common in patients with rectal CD but can occur in the absence of mucosal inflammation and is more resistant to current biological and non-biological therapies. Furthermore, fCD causes significant morbidity and psychosocial distress and is a common reason for permanent stoma formation. The pathogenesis of fCD is poorly understood, in part due to the heterogeneous clinical picture of perianal, rectovaginal and enterocutaneous phenotypes, which may each reflect a different etiology and pathophysiology. Indeed, fCD represents one of the most disabling manifestations of CD due to complete destruction of the affected mucosa, which is replaced by granulation tissue and associated with changes in tissue organization. To date, the molecular mechanisms underlying perianal fistula formation are not well defined.

Mesenchymal stem or stromal cells (MSCs) are multipotent cells capable of differentiation into a variety of cell types of mesenchymal and non-mesenchymal origin (García-Bernal et al., 2021; Harman et al., 2021; Ko et al., 2021). They can be isolated from multiple sites, most commonly adipose tissue and bone marrow. Besides their regenerative properties, MSCs have been shown to play important anti-inflammatory and immunomodulatory roles and their safety and efficacy was reported in the treatment of IBD (Dave et al., 2024; Qiu et al., 2024; Villablanca et al., 2022). Moreover, several clinical trials have reported safety and efficacy of MSCs in the treatment of perianal fCD (Ciccocioppo et al., 2011; García-Olmo et al., 2005; Molendijk et al., 2015; Panes et al., 2016; Vosough et al., 2022; Wang et al., 2023; White et al., 2024), and adipose tissue-derived MSCs (AT-MSC) have been licensed by the European Medicines Agency as a treatment for a limited phenotype of fCD (Georgiev-Hristov et al., 2018).

Although local application of human adipose tissue-derived mesenchymal stromal cells (INN Darvadstrocel; Takeda) is an approved therapy for this condition, difficulties in studying human clinical tissue and lack of an experimental model system have prevented deeper elaboration of the mechanisms underlying the therapy (Flacs et al., 2019; Li et al., 2023; Li et al., 2024). We have established a model system that recapitulates the clinical condition using human gut that has been transplanted subcutaneously in SCID mice. Fetal gut is obtained from pregnancy terminations performed legally at 12-18 weeks gestational age and transplanted subcutaneously in mature SCID mice, where it grows and can be experimentally manipulated over the course of the subsequent several months.

This experimental model system was extensively developed and refined in our laboratory for the study of human enteric nervous system (Inlender et al., 2021; Nagy et al., 2017), human-specific pathogens (Golan et al., 2011; Golan et al., 2009; Nissim-Eliraz et al., 2017) and for the development of cell-based therapies in human IBD (Canavan et al., 2016; Goldberg et al., 2019; Morris et al., 2021; Nissim-Eliraz et al., 2021). Moreover, we further refined this experimental platform for the study of fistulizing human gut disease (Bruckner et al., 2019). One of the limitations of our model system is that immune and inflammatory cellular responses are by in large mediated by host mouse cells. Nevertheless, the functionality of human-mouse interspecies systems was demonstrated extensively both in-vivo and in-vitro (Fahoum et al., 2024).

In this study, we comprehensively evaluated the preclinical safety and efficacy of human AT-MSCs in a mouse model of fistulizing gut disease. The human gut xenograft model in mice successfully replicated key features of fistulizing Crohn’s disease (fCD). Based on macroscopic observations of fistula skin wounds and histological analyses, we found that local administration of AT-MSCs was safe and significantly promoted fistula wound healing. The therapeutic effects were attributed to the prolonged survival of AT-MSCs, their active migration into the inflamed fistula tract, and their acquisition of immune-active properties, including the expression of SOD2 and CCL2, which contributed to the polarization of local macrophages into an M2 anti-inflammatory state. In contrast, AT-MSCs in healthy tissue remained stationary, adopting a fibroblast-like phenotype surrounded by extracellular matrix (ECM) and collagen, and likely did not contribute to the healing of the fistula wound.

## MATERIALS AND METHODS

### Generation and Characterization of Human Adipose-Derived MSCs

Human allogeneic expanded adipose tissue-derived mesenchymal stromal cells (AT-MSC; kindly provided by Takeda) were derived from a single healthy female donor and characterized as previously described (Lopez-Santalla et al., 2017; Lopez-Santalla et al., 2018; Mancheño-Corvo et al., 2017). Human samples were obtained after informed consent as approved by the Spanish Ethics Committee of reference for the site of tissue procurement (Clinica de la Luz Hospital, Madrid, Spain). Cells were expanded up to duplication 12–14 and frozen. AT-MSC were thawed, seeded in tissue culture flasks, and trypsinized before administration to mice. AT-MSC were defined according to the criteria of the International Society for Cellular Therapy, being positive for CD73, CD90, and CD105 and negative for CD14, CD19, HLA-DR, CD34 and CD45 as previously described (Dominici et al., 2006).

Lentivirus-based transduction systems was used as fluorescence reporting systems that will enable visualization and isolation of AT-MSC in tissues. FUGW (a gift from David Baltimore to Addgene plasmid # 14883; http://n2t.net/addgene:14883; RRID:Addgene_14883) (Lois et al., 2002) was used as target plasmids (transfer vectors) constructed to constitutively express eGFP in AT-MSc with lentivirus constructs as previously reported by us (Nissim-Eliraz et al., 2021; Salamon et al., 2020). For lentivirus preparation HEK 293-FT cells were grown in high glucose DMEM with 0.1mg/ml Pen-Strep, 2mM L-glutamine, 0.1mM Non-Essential Amino Acids (NEAA), 1mM Sodium Pyruvate and 10% fetal calf serum (FCS). Cells were seeded at 1x10^7^ cells/plate using 150 mm plates, precoated with Poly-l-lysine (Sigma P1274). The next day each plate was transfected using PEI (Polysciences) with 20ug of a mixture containing the lentiviral vector plasmid and FUGW with pMDLg/RRE packaging plasmid, pRSV-Rev plasmid and pCMV-VSV-G at 4:2:2:1 ratio respectively. The next morning cells were washed twice with PBSX1 and the medium was replaced to growth media with 2%FCS. Viral particles were harvested two days later by collecting the medium from each plate and filtering through 0.45um filter. Virus was concentrated by ultracentrifugation at 4° C, 79,000 g for 2 hours (sw28 rotor). The pellet from each tube was resuspended in 120ul sterile PBSX1, aliquoted and frozen at -80° C. For transduction of AT-MSC, cells were plated on six-well plates at a concentration of 15x10^4^ cells/well in MSCs media (high glucose DMEM with 0.1mg/ml Pen-Strep, 2mM L-Alanine-L-glutamine and 10% fetal bovine serum). Twenty microliters FUGW lentiviruses were directly overlaid on cells and incubated for 3 days. Cells were then washed, trypsinized and transferred to 10ml plate with fresh media. Five days later the cells were harvested by trypsinization, washed and spinned for 1200rpm for 5 min. Fluorescent intensity was analyzed microscopically and by NovoCyte Quanteon Flow Cytometer (Agilent) with 10,000 cells per sample. The cells were gated for GFP signals based on the background signal from the non-transformed cells. The fraction of fluorescent AT-MSC cells following transduction with 20 μl of lentivirus FUGW was 99.3% (Supplementary figure 1).

### SCID mouse human intestinal xeno-transplant model

C.B-17/IcrHsd-Prkdcscid (abbreviated as SCID) mice were purchased from Envigo Israel (Rehovot, Israel). All mice were housed in a pathogen-free facility, in individually ventilated cages (IVC), given autoclaved food and water. All animal use was in accordance with the guidelines and approval of the Animal Care and Use Committee (IACUC) of the Hebrew University of Jerusalem. IRB and IACUC approvals were obtained prospectively (Ethics Committee for Animal Experimentation, Hebrew University of Jerusalem; MD-18-15502-5 and MD-20-16275-5 and the Helsinki Committee of the Hadassah University Hospital; 81-23/04/04). Women undergoing legal terminations of pregnancy gave written, informed consent for use of fetal tissue in this study. Human fetal small bowel 12-18 weeks gestational age was implanted subcutaneously on the dorsum of the mouse as described previously (Bruckner et al., 2019; Canavan et al., 2016; Golan et al., 2011; Golan et al., 2009; Goldberg et al., 2019; Nagy et al., 2017; Nissim-Eliraz et al., 2021; Nissim-Eliraz et al., 2017). All surgical procedures were performed in an aseptic working environment in a laminar flow HEPA-filtered hood with isoflurane inhalation anesthesia (1.5 to 2% v/v isofluorane in O2). Before surgery, carprofen (5 mg/kg, Rimadyl, Pfizer Animal health) was administered subcutaneously. The surgical area was shaved and depilated (Nair hair removal cream) and the skin was scrubbed and disinfected with betadine and 70% (v/v) ethanol. After surgery the mice were provided with enrofloxacin-medicated water (Bayer Animal HealthCare AG) for 7 days and were closely monitored once a day for behavior, reactivity, appearance and defecation. Grafts developed in situ for 12-16 weeks prior to manipulation.

### Fistula model system

Enterocutaneous fistulas were induced in female SCID mice with fully developed human gut xenografts (12-16 weeks after transplantation) by combining Imiquimod-induced psoriatic dermatitis (van der Fits et al., 2009) with systemic LPS treatment (Morris et al., 2021). Mice were depilated 2-3 days prior to the commencement of the experiment. To this end, mice were anesthetized with 2% isoflurane in oxygen, and all hair removal took place under a hood. A rectangular region of hair on the dorsal side, from the hind legs to the front legs, was first removed using an electric razor (Norelco G390, Philips), and the skin was then depilated using the chemical Nair® (Church & Dwight Co., Inc.). Next, mice received a daily topical dose of 62.5 mg of 5% Imiquimod (IMQ) cream (3.125 mg of the active compound) on the shaved and depilated skin overlying the transplants for 7 consecutive days. On the fourth day, mice received a single treatment of 0.1 mg/kg LPS by IP injection as previously described (Morris et al., 2021).

This treatment protocol was empirically determined to cause most optimal and reproducible fistulating gut xenografts in the model system. Fistulas developed in approximately 50% of human gut xenografts within 7 to 21 days following commencement of the induction protocol. The utility of the IMQ protocol for the induction of psoriasis-like disease in SCID mice used in our model system was initially verified and results are presented in the Supplementary materials (Supplementary Figure 2).

### AT-MSC treatment

In vitro cultured AT-MSCs (1.0 × 10^6^) were resuspended in 50 µl PBS and injected subcutaneously around the fistula skin wound. Animals were randomly assigned into AT-MSC treatment group (n=5) or control untreated group (n=5). Fistula wound and fistula tract did not undergo debridement or curettage and injection area was not scrubbed or disinfected. Mice did not receive any antimicrobial or anti-inflammatory treatment in the whole duration of the experiment. Control untreated mice did not receive shame treatment of PBS injection considering the risk of introducing and spreading infection and exacerbating wound condition.

Immediately before treatment and thereafter once weekly images were taken of the wound and a ruler at level for scale. Using QuPath (Bankhead et al., 2017) and Fiji (Schindelin et al., 2012), the circumference of the wounds in each image was traced and the area was determined. Data were plotted as percentages of the initial wound area (100%).

All animals were sacrificed 3 weeks after a single AT-MSC treatment or no-treatment protocol. Closure of the wounds was then monitored daily by measuring the diameter and circumference. Images of scaled wounds were digitally analyzed and wound areas and circumference were measured using QuPath and Fiji softwares as previously described (Yampolsky et al., 2024).

### Tissue collection and analysis

From each AT-MSC-treated and untreated control mouse, dissection was performed to in situ encompass marginal normal skin, fistula wound, underlying fistula tract and human gut xenograft to preserve the anatomical structures and spatial arrangement of the enterocutaneous fistulizing human gut. Tissues were FFPE and OCT-embedded for cryo blocks and sectioned for H&E and immune staining as previously described (Schneider et al., 2023). All Bright-field microscopic Images were acquired using a 3D HISTECH Pannoramic-250 microscope slide-scanner (3D HISTECH, Budapest, Hungary). Snapshots were taken with Case Viewer software (3D HISTECH, Budapest, Hungary). All quantitative analysis; thickness measurements, collagen staining and CD45 staining, were performed using ImageJ software package.

Sequential tissue sections were analyzed using H&E and immunofluorescence staining for microscopy and for spatially-resolved whole-transcriptome analysis by Nanostring GeoMx digital spatial profiling (DSP).

Histological quantification for each wound bed was conducted on the two central-most sections from each animal. Tissue sections were harvested (fixed in formalin), processed (embedded in paraffin and sectioned at 5 μm), deparaffinized (xylene immersion), and stained (H&E, MT) using standardized histological procedures. The percentage of the wound bed covered by re-epithelialization was calculated from the 2 central most tissue sections using ImageJ software (National Institutes of Health, Bethesda, MD) as described previously (Fuchs et al., 2021; van de Vyver et al., 2021).

### Immunohistochemistry

FFPE tissues and OCT-embedded cryo sections were stained for epifluorescence and confocal microscopy as previously described (Schneider et al., 2023). Incubation with primary antibodies included: rabbit anti-human CD45 (1:25) (Abcam, Cambridge, UK), rat anti-mouse CD45 (1:50) (Abcam, Cambridge, UK), rabbit anti-GFP (1:100) (Abcam, Cambridge, UK), or rabbit anti-human/mouse SOD2 (Proteintech, Rosemont, Illinois, USA) with 2% normal donkey serum (NDS), overnight at 4 °C. Secondary antibody labelling included: Alexa Fluor 488 donkey anti-rat, Alexa Fluor 594 or 647 donkey anti-rabbit, Alexa Fluor 488 donkey anti-mouse (all 1:500; invitrogen Waltham, Massachusetts, USA) with 2% NDS for 1 h at room temperature. DAPI was used for nuclear staining and F-actin was stained in frozen sections using phalloidin.

Microscopic analysis was performed using M1 Imager Axio epifluorescence microscope with an MRm Axio camera (Zeiss), and images were captured using Zen software V3.4. Confocal microscopy was performed using Nikon Ti2E fitted with a Yokogawa W1 Spinning Disk and images were captured and processed for publication using NIS-Elements Package.

### Spatial Transcriptomics

Duplicate sequential formalin-fixed paraffin-embedded (FFPE) 5 μm sections of AT-MSC-treated (n-3) and non-treated fistula tissues (n=3) were prepared to NanoString specifications on Lecia Apex BOND Superior Adhesive slides (3800040). Slide were sent to GeoMx DSP (digital spatial profiling) whole transcriptome atlas (WTA) analysis (Nanostring Technology Access Program, Seattle, USA). Duplicate sequential sections were required to separately analyze gene expression in AT-MSC and in mouse cells constituting the fistula tract.

Tissue slides were baked in a drying oven at 60 °C for 1 h and then deparaffinized and rehydrated using standard procedures. After the target retrieval step, tissues were treated with proteinase K solution to expose RNA targets followed by fixation with buffered formalin. After all tissue pretreatments were done, tissue slides were then stained using nuclear stain Syto83 (Thermo, S11364), anti-GFP (Thermo, A21311) to visualize AT-MSC and anti-PanCK (Novus, NBP2-33200AF488, clone # AE1/AE3) for tissue visualization, and incubated with RNA probe mix (in situ hybridization probes with UV photocleavable oligonucleotide barcodes, GeoMX human or mouse Whole Transcriptome Atlas kit). Regions-of-interest (ROI) were subsequently selected on the basis of the distinct fluorescent signals from PanCK and GFP staining designated as MSC-positive and MSC-negative fistula ROIs, and perifistula and normal ROIs. The UV cleavable oligo tags were then collected from the designated regions and separately deposited into 96-well plate. The resulting mRNA samples were subjected to sequencing using Illumina’s NovoSeq instrument. Raw fastq files along with the configuration file were used as input to the GeoMx ngs pipeline command line tool running on a Linux server. We used NanoString’s GeomxTools Bioconductor package (https://www.bioconductor.org/packages/release/bioc/html/GeomxTools.html) for downstream data analyses. These included QC & preprocessing, data normalization, unsupervised clustering, and differential gene expression. For segments that passed these QC parameters, we utilized Q3 (3^rd^ quartile of all selected targets) normalization and mixed effect modeling for identifying differentially expressed genes as presented in the results.

### Pre-ranked GSEA analysis

Genes were ranked based on t statistics from differential gene expression analysis. Pre-ranked GSEA analysis was then conducted using the GSEA software (GSEA 4.1.0) (Mootha et al., 2003). Genes were mapped to gene ontology biological processes pathways obtained from the Molecular Signatures Database (MSigDB) (Liberzon et al., 2011). The results of pathway analysis were then visualised using the ggplot2 and ComplexHeatmap package in R. Dotplots presented Normalized Enrichment Score (NES), FDR adjusted p-values as -Log10 and the Enrichment Signal Strength (ESS) as explained in https://www.gsea-msigdb.org/gsea/doc/GSEAUserGuideFrame.html. GeoMx gene expression data of AT-MSC was further analyzed for cell classification using xCell, a computational algorithm published in the journal, Genome Biology in 2017 by Aran et al (Aran et al., 2017).

### Statistical analysis

Statistical analyses of data, unless otherwise stated, were conducted using GraphPad Prism (10.1.2) software. Specific statistical tests used for each dataset and error bars (SD or SEM) are stated in respective figure legends. P < 0.05 was considered significant.

## RESULTS

### Locally administered AT-MSC improved fistula skin wound healing

After completing the IMQ-LPS induction protocol, treated and untreated fistula skin wounds were monitored for 3 weeks, covering the initial stages of normal wound healing (van de Vyver et al., 2021). Measurements of skin wound area and perimeter showed that local application of AT-MSC improved fistula wound healing compared to untreated controls (Figure 1A-F). Tissues were then collected in situ, and healing dynamics were assessed histologically, focusing on re-epithelialization, epithelial thickness, keratinization, and scab thickness. Skin wound re-epithelialization was significantly enhanced in the AT-MSC treated group (Figure 1G-I). Additionally, the wound scab remained attached to the epidermis in control mice, a phenomenon linked to poor re-epithelialization.

**Figure 1.**
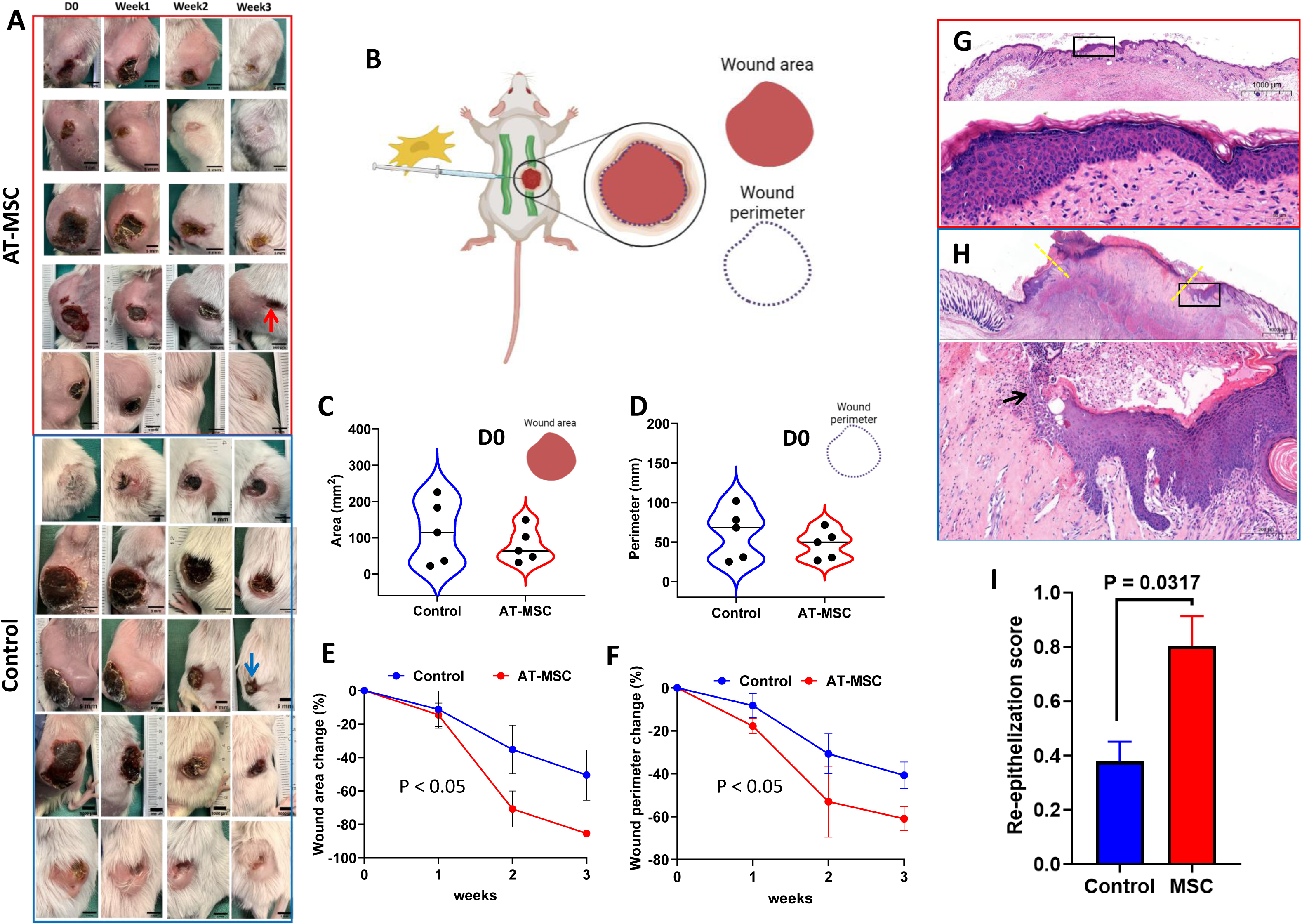
Locally administered AT-MSC enhanced fistula skin wound healing by expediting wound re-epithelization. Enterocutaneous fistulas developed in SCID mice transplanted with human gut xenografts using the IMQ-LPS induction protocol. Fistula skin wounds were imaged at treatment day and at least once weekly thereafter (A). AT-MSC (10^6^ cells in 50µl saline) were injected around the fistula tract at D0 (B). Wounds’ area and perimeter were measured using scaled images in Fiji software, and no significant difference was found between the AT-MSC-treated and control groups at D0 (C-D). The efficacy of AT-MSC therapy is supported by the visual appearance (right raw in A) and measured areas and perimeters of fistula wounds after 3 weeks (E and F, respectively). Data is presented as mean ± SEM and significance of comparison between treatment groups was determined by one-way ANOVA and Dunnett’s test. Representative histological images of fistula skin wounds harvested 3 weeks after treatment with AT-MSC (G) and Non-treated control (H). FFPE sections of two central-most sections were stained with H&E and re-epithelization was measured and presented as fraction of epithelized wound (C). Boxed areas in top panels in G-H are shown in higher magnification in bottom panels, yellow lines indicate the wound margin, and black arrow indicates the front of the neo-epidermis. Note the attachment of a scab to the wounded epidermis in control mice. Scale bars; 50µm (G; bottom panel), 200µm (H; bottom panel) and 1000µm (G and H; top panels). Images were acquired using a 3D HISTECH Pannoramic-250 microscope slide-scanner (3D HISTECH, Budapest, Hungary). Snapshots were taken with Case Viewer software (3D HISTECH, Budapest, Hungary). Measurements of wound size and epithelization were performed using QuPath software. Error bars indicate mean ± SEM and statistical analysis was performed using non-parametric t-test. All graphs and statistical analyses were performed in Prism, GraphPad and P < 0.05 was considered statistically significant. Illustration B was created with BioRender.

### Administered AT-MSC formed two distinct cell populations: reactive and stationary

AT-MSC expressing eGFP were detected in treated fistulas 3 weeks after injection through fluorescence microscopy (Figure 2A-H). These cells appeared in two distinct locations: along the fistula tract and in the surrounding normal skin. The morphology and arrangement suggested that AT-MSC within the fistula tract were migratory and reactive (Figure 2G-H), while those in the normal skin were stationary, compacted in the extracellular matrix, and enclosed by a fibrous capsule (Figure 2B-E and schematically summarized in Figure 2J). FFPE sections were stained with SYTO88 for nuclear visualization, anti-pan-cytokeratin, and anti-GFP antibody. Using GeoMx DSP, a whole transcriptomic analysis was conducted on the described AT-MSC populations in the tissues. Additionally, the same AT-MSC used for treatment were cultured on chamber slides (µ-Slide 8 well; ibidi), stained with SYTO88 and anti-vimentin antibody, and analyzed alongside the FFPE samples. Principal Component Analysis (PCA) using GeoMx DSP expression data from all cell samples demonstrated alignment of the described cell phenotypes (Figure 2K). Gene expression between the two in vivo phenotypes—reactive and stationary—and the common origin of all cells, the in vitro cultured cells, was compared. The analysis is presented through Venn diagrams (Figure 3A-B), volcano plots, and MA plots (Figure 3C-D for reactive vs in vitro, and Figure 3F-G for stationary vs in vitro). Differentially expressed genes were further examined using GSEA, revealing the immune activity of reactive cells (Figure 3E) and the fibroblast-like activity of stationary cells (Figure 3H). Additionally, gene expression in reactive AT-MSC versus stationary AT-MSC was compared using volcano and MA plots (Figure 4A-B). Notably, the mitochondrial enzyme superoxide dismutase (SOD2) and chemokine CCL2 were overexpressed in reactive AT-MSC located in the fistula tract, which was validated by immunofluorescence staining for SOD2 (Figure 4F-G). Furthermore, cell classification via xCell (Figure 4C), GSEA (Figure 4D), and STRING pathway analysis (Figure 4E) clearly indicated that stationary AT-MSC had changed into fibroblast-like cells, whereas reactive cells in the fistula tract had developed into immune-active cells.

**Figure 2.**
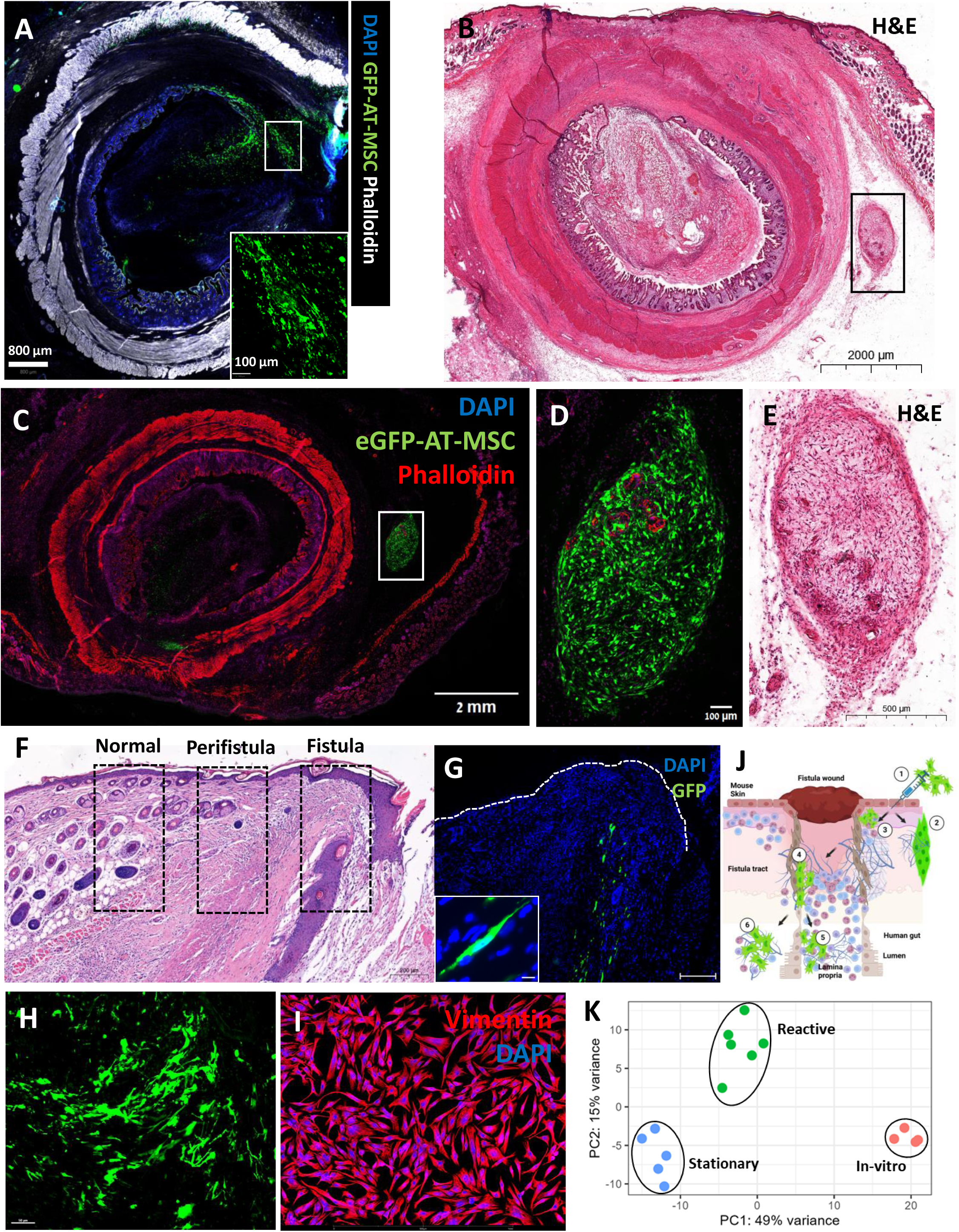
We show here that AT-MSC cultured in-vitro and locally administered around the fistula wound survived for 3 weeks in vivo. AT-MSC expressing GFP were directly visualized (A and inset A, C, and H), or indirectly using anti-GFP antibody (G and inset G, dotted line indicates outer border of epidermis), by epifluorescence microscopy. AT-MSC were characterized into 2 phenotypically distinct population designated as “stationary” which were located in normal skin areas (boxed areas in B-C, shown in higher magnification in D-E, respectively) and “reactive” which were located in the fistula tract. Schematic illustration (created with BioRender) summarizing in-vivo traffic routes of AT-MSC transplanted around the fistula wound (J). GeoMx DSP was used to concurrently analyze the transcriptome of stationary (C-D) and reactive (G-H) AT-MSC in FFPE sections and a sample of the treatment AT-MSC grown on glass slide (I). PCA analysis of GeoMx DSP data demonstrates clustering of samples and phenotypes (K). Scale bars; 800 µm (A), 2000 µm (B-C), 100 µm (D and inset A), 500 µm (E), 200 µm (F), 50 µm (G-I), and 10 µm (inset G).

**Figure 3.**
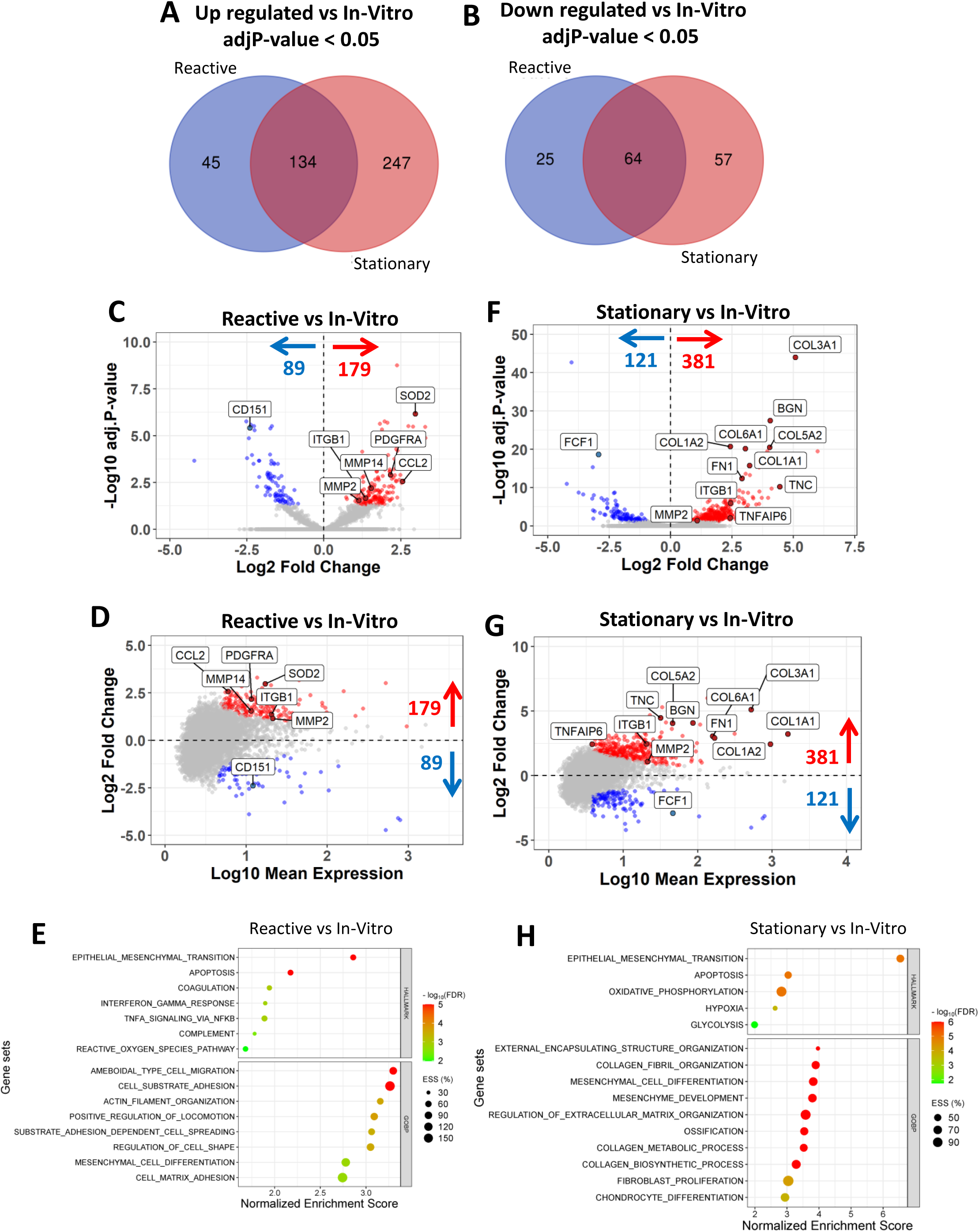
We show here that AT-MSC, which originated in vitro and were administered around the fistula wound, exhibit different cellular characteristics based on their location, as described in the Figure 2. GeoMx DSP was employed to concurrently analyze the transcriptomes of stationary and reactive AT-MSC in formalin-fixed paraffin-embedded (FFPE) tissue sections obtained from treated fistulas, as well as AT-MSC cultured on glass slides in vitro. Gene expression comparisons between reactive AT-MSC located in the fistula tract and in vitro cells are presented using Venn diagrams (A-B), a volcano plot (C), and an MA plot (D). Similarly, gene expression comparisons between stationary AT-MSC in normal tissue and in vitro cells are shown using Venn diagrams (A-B), a volcano plot (F), and an MA plot (G). Gene Set Enrichment Analysis (GSEA) of the expression data indicates that reactive cells in the fistula tract exhibit immune activity, while stationary cells display a fibroblast-like phenotype. Pre-ranked GSEA analysis was conducted using GSEA software (GSEA 4.1.0).

**Figure 4.**
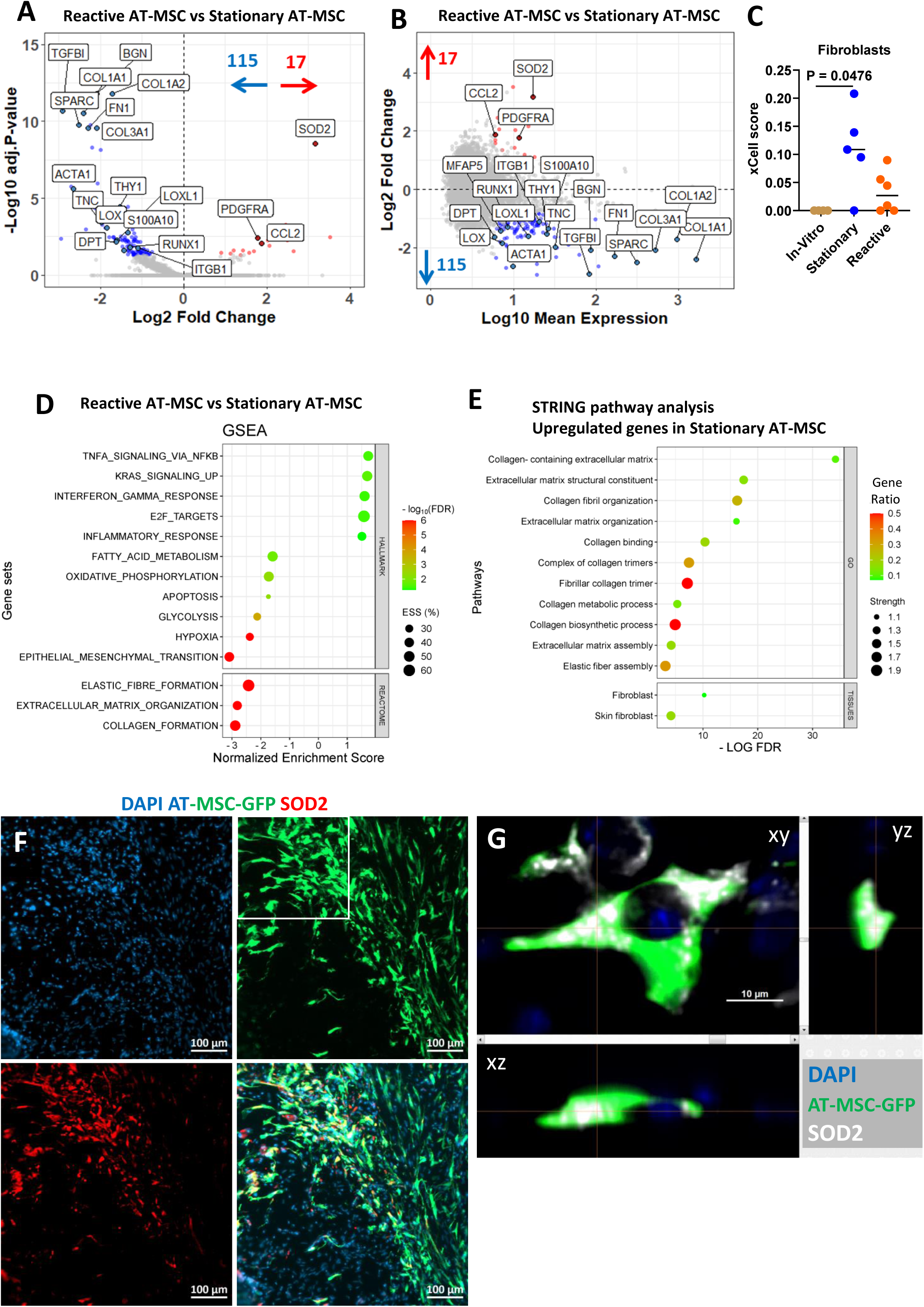
AT-MSC transplanted around the fistula skin wound adopted immune reactive phenotype in the fistula tract and stationary fibroblast-like cells in the normal tissue. GeoMx DSP was used to concurrently analyze the transcriptome of stationary and reactive AT-MSC in FFPE tissue sections obtained from 3 weeks after transplantation in the tissues around the fistula wound. GeoMx gene expression data of AT-MSC was analyzed for cell classification using xCell (C). Pre-ranked GSEA analysis was conducted using the GSEA software (GSEA 4.1.0) and results are presented in dot plot D. Positive normalized enrichment scores (NES) are enriched in reactive AT-MSC (D) while negative NES are enriched in stationary AT-MSC (D). Pathway enrichment analysis using STRING was applied to all significantly downregulated genes (blue data points in A-B), which are up regulated in stationary cells (n=115; adjP-Value < 0.05) and results are presented in dot plot E. Expression of SOD2 in AT-MSC is further validated using immunofluorescence staining in F-G. Representative images of AT-MSC in the fistula tract expressing eGFP in green, nuclear staining with DAPI in blue and anti-SOD2 in red. Colocalization of SOD2 (white in G) and GFP (green in G) in AT-MSC was demonstrated using confocal microscopy. The xy image is on the plan indicated by the horizontal and vertical lines shown in the xz and yz images, respectively, and the original magnification was X40 (G). Scale bars: 100µm (F), and 10µm in G.

### Reactive AT-MSC created a niche of anti-inflammatory macrophages in the fistula tract

We have previously demonstrated the presence of resident human immune cells in the mature gut xenograft (Bruckner et al., 2019). However, the immune cell response in the fistula tract is entirely murine, while human immune cells which are present in the fistulated human gut, are not present in the fistula tract (Supplementary Figure 3). This notion is further supported by the lack of expression of human PTPRC gene (CD45) in the fistula tract regions analyzed by GeoMx DSP as described above (Figure 2). To this end, we next used the GeoMx DSP system to analyze murine gene expression in the previously defined regions (designated as normal, perifistula and fistula regions in Figure 2F) on the sequential FFPE tissue sections. The fistula region was further designated as histologically positive for the presence of AT-MSC (MSC+ve based on visualized GFP) and histologically devoid of AT-MSC (MSC-ve), which was further validate by the huge difference in EGFP gene expression between the above predefined regions (Figure 5A and 5E). PCA was performed using the GeoMx DSP expression data of all of these samples showing alignment of the above described samples and sub-regions (Figure 5B). Differential gene expression between MSC-ve fistula regions, MSC+ve fistula region, and perifistula regions versus the normal regions is presented using Venn diagram (Figure 5C), volcano plots (Figure 5D-F, respectively) and MA plots (Figure 5D-F, respectively). GSEA analysis of the differentially expressed genes shows significant enrichment of inflammation hallmark gene sets in the fistula and perifistula regions (see insets of mountain plots in right panels of Figure 5D-F) and significant enrichment of GOBP macrophage activation gene set in the MSC+ve region (see inset of mountain plot in left panel of Figure 5E). To better understand the effects of AT-MSC on the inflamed fistula we next compared the gene expression profiles between the MSC+ve and MSC-ve fistula regions (Figure 6). This analysis revealed significant upregulation of many macrophage marker genes (Garrido-Trigo et al., 2023; Mulder et al., 2021) in the MSC+ve regions such as Lyz2, Adgre1 (F4/80), Cd68, Aif1 (IBA-1), Cd14, C1qa, C1qb, and Itgam (Cd11b) (Figure 6), further supporting the notion that MSC+ve regions are enriched with macrophages. GSEA of MSC+ve versus MSC-ve expression data based on the ImmuneSigDB gene set collection (MSigDB, Broad institute) revealed highly significant enrichment of M2-macrophage gene set (Figure 6D-E; based on (Edwards et al., 2006)) in MSC+ve regions and M1-macrophage gene set in MSC-ve regions (Figure 6D&F; based on (Szanto et al., 2010)). Specifically, the M2-genes Cd163, F13a1, Retnla, and Apoe, Cd209a, Cd209b, Cd86, Trem2, and Ccl1 were significantly overexpressed in the MSC+ve regions. This was further validated through fluorescent immune staining of control, non-treated fistula and AT-MSC treated fistula tissues using antibodies to the macrophage marker IBA-1 and M2 marker CD163 (Figure 6G-H, Supplementary Figure 4 and Supplementary Figure 5). Moreover, many other gene known to be associated with M2-polarization were also significantly upregulated these include Stab1, Fn1, Alox5, Irf4, Pparg, Mdk, and Fpr2 (Figure 6).

**Figure 5.**
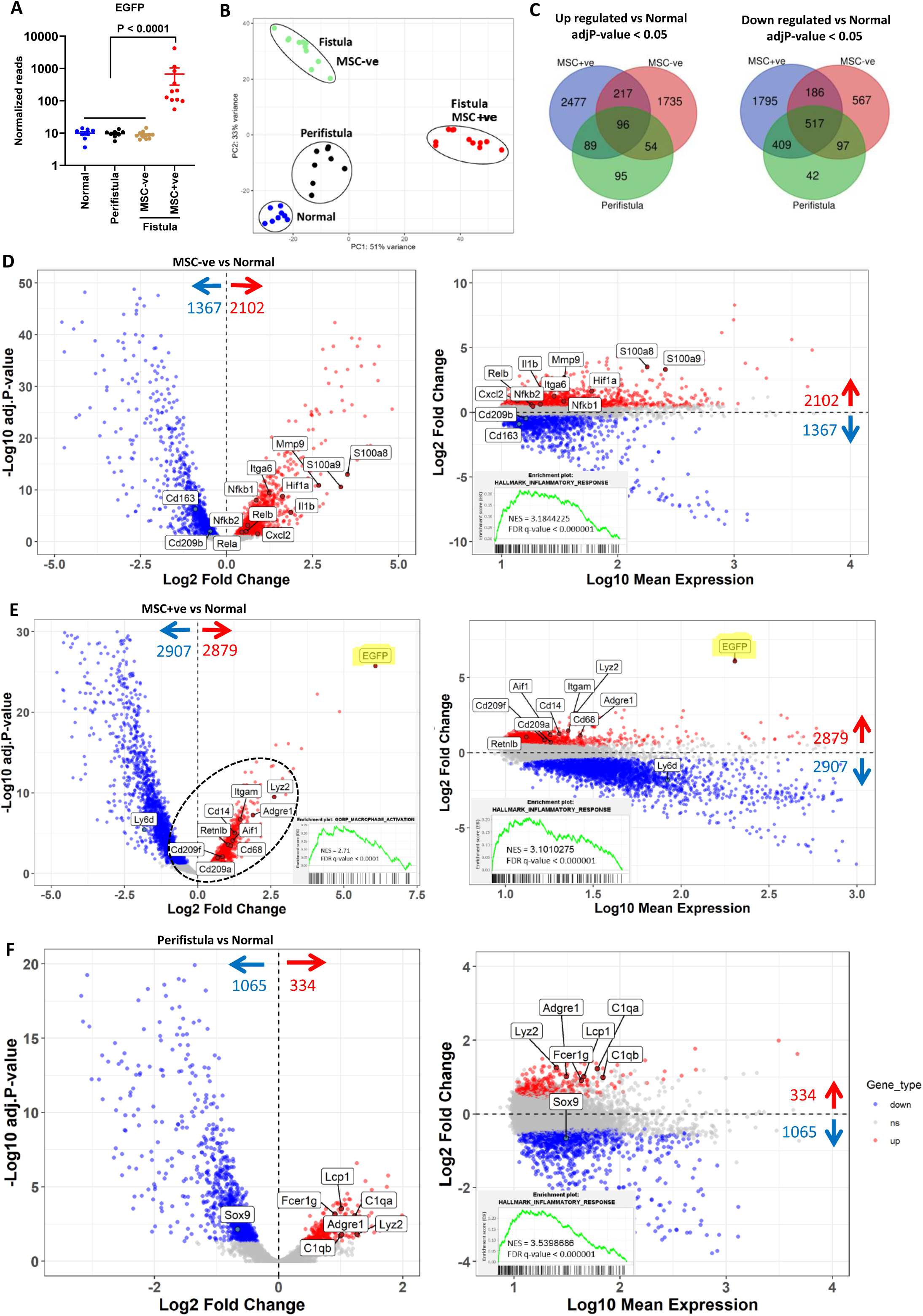
Spatial transcriptomic analysis of treated and non-treated control fistulas using GeoMx DSP. The predefined fistula regions (Figure 2F) were analyzed for murine transcriptomic expression and the fistula regions were further divided to MSC+ve and MSC-ve regions based on fluoresnce microscopy which was further validated based on EGFP gene expression (A). GeoMx DSP data was analyzed using PCA showing alignment of samples and sub-regions (B). Differential gene expression comparing MSC-ve fistula regions, MSC+ve fistula region, and perifistula regions versus the normal regions is presented using Venn diagram (C), volcano plots (D-F, respectively) and MA plots (D-F, respectively). GSEA analysis of differentially expressed genes was performed based on the Hallmark gene set showing significant enrichment for inflammation by mountain plots, normalized enrichment score (NES) and adjusted P-value (insets in MA plots D-F). MSC+ve region also showed enrichment of GOBP macrophage activation gene set (mountain plot, NES and adjusted P-value (inset in volcano plot E).

**Figure 6.**
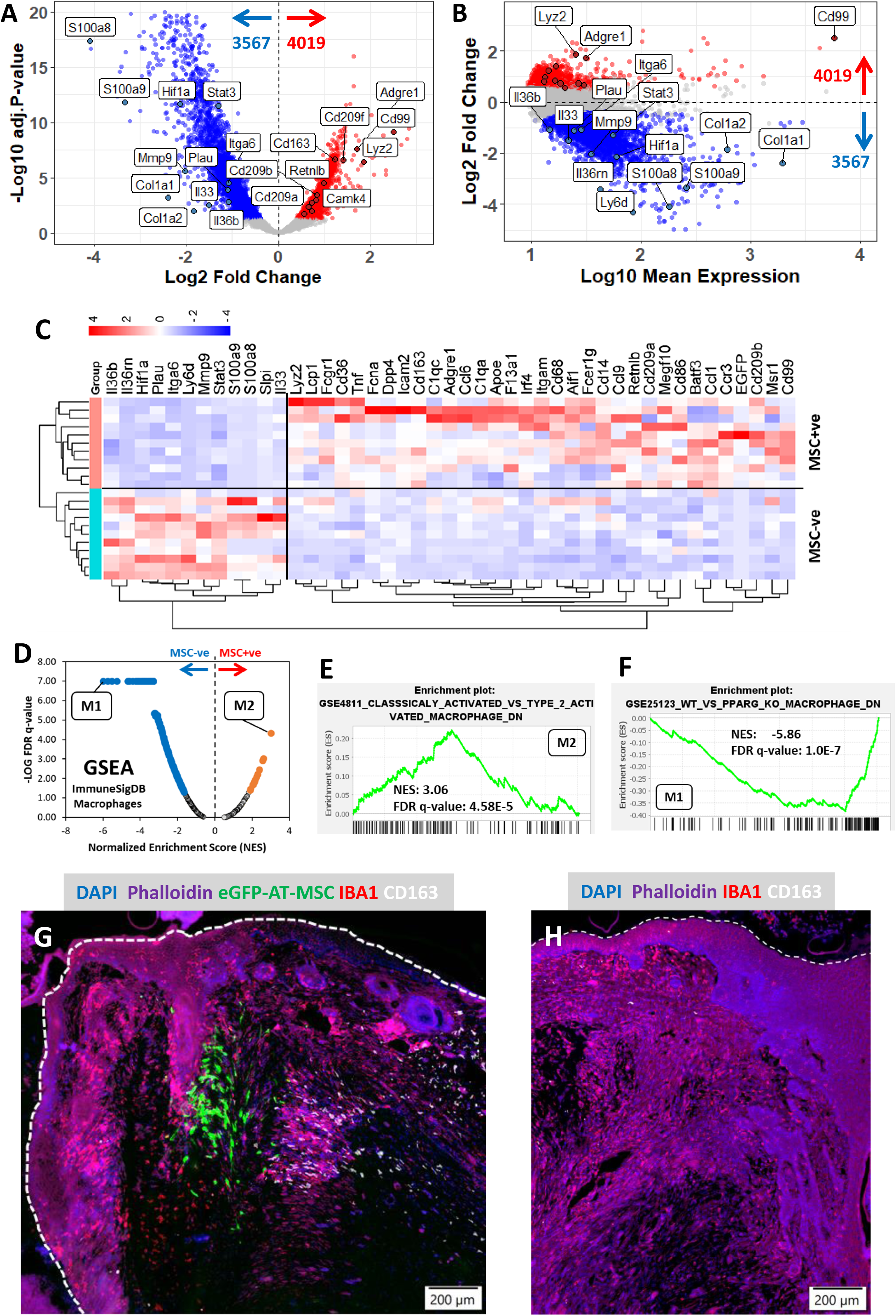
MSC+ve region is enriched with M2 macrophages. GeoMx DSP expression data was analyzed for differentially expressed genes (DEG) in MSC+ve versus MSC –ve regions and data is presented as volcano plot (A) and MA plot (B) highlighting many macrophage markers genes over expressed in MSC+ve regions. Selected genes are presented in heatmap C (created with Srplot). Gene set enrichment analysis (GSEA) was performed using the ImmuneSigDB gene set collection (4872 gene sets) comparing MSC+ve regions to MSC-ve regions. All GSEA results containing macrophage data are visualized by a vulcano plot (D) representing the normalized enrichment scores (NES) on the x-axes and the FDR-adjusted (False-discovery-rate) p values (-log10) on the y-axes. Red dots and blue dots in D represent significantly enriched gene sets in the MSC+ve regions and MSC-ve regions, respectively, with FDR adjusted P-value < 0.05. Selected top ranking gene sets are highlighted in the text box and further presented in the mountain plots E and F. Enrichment of M2 macrophages in AT-MSC (eGFP in G) treated fistula tracts was further validated using fluorescence microscopy of tissue sections stained with DAPI (blue in G), phalloidin (pink in G), antibodies to IBA1 (red in G) and CD163 (white in G). Non-treated fistula was similarly stained demonstrating dearth of CD163-positive macrophages in the fistula tract (H). Individual channels of G and H are shown in Supplementary Figure 4 and supplementary Figure 5, respectively. Image was acquired using Olympus SLIDEVIEW VS200 scanner and snapshots were taken with OlyVIA v. 4.1.1 software. Scale bar; 200µm (G-H).

## DISCUSSION

In this study, we demonstrated the safety and efficacy of clinical-grade adipose-derived human mesenchymal stromal cells (AT-MSC) for treating gut fistulizing disease. We also explored several aspects of the mode of action of these cells in vivo, which may be relevant for clinical applications in patients. Crohn’s disease (CD)-associated enterocutaneous fistulizing disease is currently a chronic, incurable condition and represents a significant clinical challenge that remains unmet. Preclinical studies primarily utilize mouse model systems. While many useful inflammatory bowel disease (IBD) models have been developed, none have effectively modelled fistulizing gut disease. To address this gap, we established a novel model of fistulizing disease using human gut transplants in mice. Our findings indicate that concurrent inflammation in the human gut transplants and the overlying skin leads to the development of fistulizing disease. Skin inflammation was induced using the topical application of imiquimod (IMQ), a widely used model for psoriatic-like disease in mice (van der Fits et al., 2009). At the same time, systemic application of lipopolysaccharide (LPS)—an essential component of gut bacterial cell walls—activated inflammation in the adjacent human gut transplants (Morris et al., 2021; Nissim-Eliraz et al., 2021; Ohishi et al., 2024). The resulting fistula skin wounds formed tracts connecting the human gut transplants and mouse skin. While clinical material, particularly following AT-MSC treatment, is not available, here we were able to thoroughly harvest the fistula skin wound, fistula tract, and portions of the human gut for in-depth histological analysis. These analyses included histopathology, high-plex immunostaining, and spatial transcriptomics of both human and mouse cells. While live MSC are rarely observed following in vivo treatment, we demonstrated that these clinically used cells survived in the model system for at least three weeks. Moreover, the cells differentiated into two distinct populations that we designated as reactive and stationary. This observation suggests that the cellular and molecular microenvironment surrounding the administered MSC creates a niche that shapes their phenotype.

Using spatial transcriptomic analysis, we showed that AT-MSC in the fistula tract exhibited an inflammatory phenotype, characterized by the overexpression of SOD2 and CCL2, whereas those in normal tissue displayed a fibroblastic phenotype. Numerous previously published in vitro and in vivo studies have highlighted the roles of SOD2 and CCL2 in the therapeutic activity of MSC (Giri et al., 2020; Lee et al., 2020; Marx et al., 2021; Rafei et al., 2008; Whelan et al., 2020). Enhanced activity has been reported with pre-activation or genetic overexpression, further validating our novel in vivo observations. Suggested mechanisms include M2 polarization of macrophages, which exerts anti-inflammatory effects. In agreement with this prior knowledge, we found that the microenvironment of the reactive AT-MSC in the fistula tract was enriched with M2 macrophages. This mechanism of action was recently suggested for rat adipose-derived stem cells (Li et al., 2023) and extracellular vesicles derived from rat mesenchymal stem cells (Li et al., 2024) using a rat model of perianal fistula (Flacs et al., 2019).

Taken together, we propose that AT-MSC injected in the periphery of the fistula wound migrate into the fistula tract, adopting an inflammatory phenotype characterized by SOD2 and CCL2 expression, enabling them to promote the polarization of recruited inflammatory M1 macrophages into M2 anti-inflammatory macrophages. The second population of AT-MSC was found to be stationary in normal tissue, encapsulated by fibrous tissue, and embedded in the extracellular matrix (ECM), with a lack of visible immune or inflammatory response. Based on our results, we suggest the following implications: (1) the IMQ-LPS model system using human gut xenografts can serve as a reliable preclinical model for fistulizing Crohn’s disease (fCD); (2) AT-MSC therapy is effective for treating fCD; and (3) the efficacy of this therapy can be further enhanced by pre-activation or genetic manipulation to boost the expression of SOD2 and CCL2.

## Acknowledgments

The authors acknowledge the technical assistance of David Scoville, Anna Pavenko and Yan Liang, Nanostring Technology Access Program (TAP), Seattle, USA, for the performance of GeoMx DSP analysis.

## Author contribution

N.Y.S conceived the study and planned experiments; performed experiments, data acquisition, and analysis and interpretation; and wrote the manuscript with input from all authors. N.M. performed experiments, sample collection, histology and spatial transcriptomics with assistance from Y.L., I.S. and E.N.E. P.S. contributed to spatial transcriptomic data analysis. S.Y. provided human fetal gut specimens. K.K. and E.L. contributed to data interpretation and manuscript review and editing.

## Data Availability Statement

All data associated with this study are present in the paper or the Supplementary Materials. The spatial transcriptomic data that support the findings of this study are available from the corresponding author upon reasonable request.

---

**Supplementary Figure 1.**
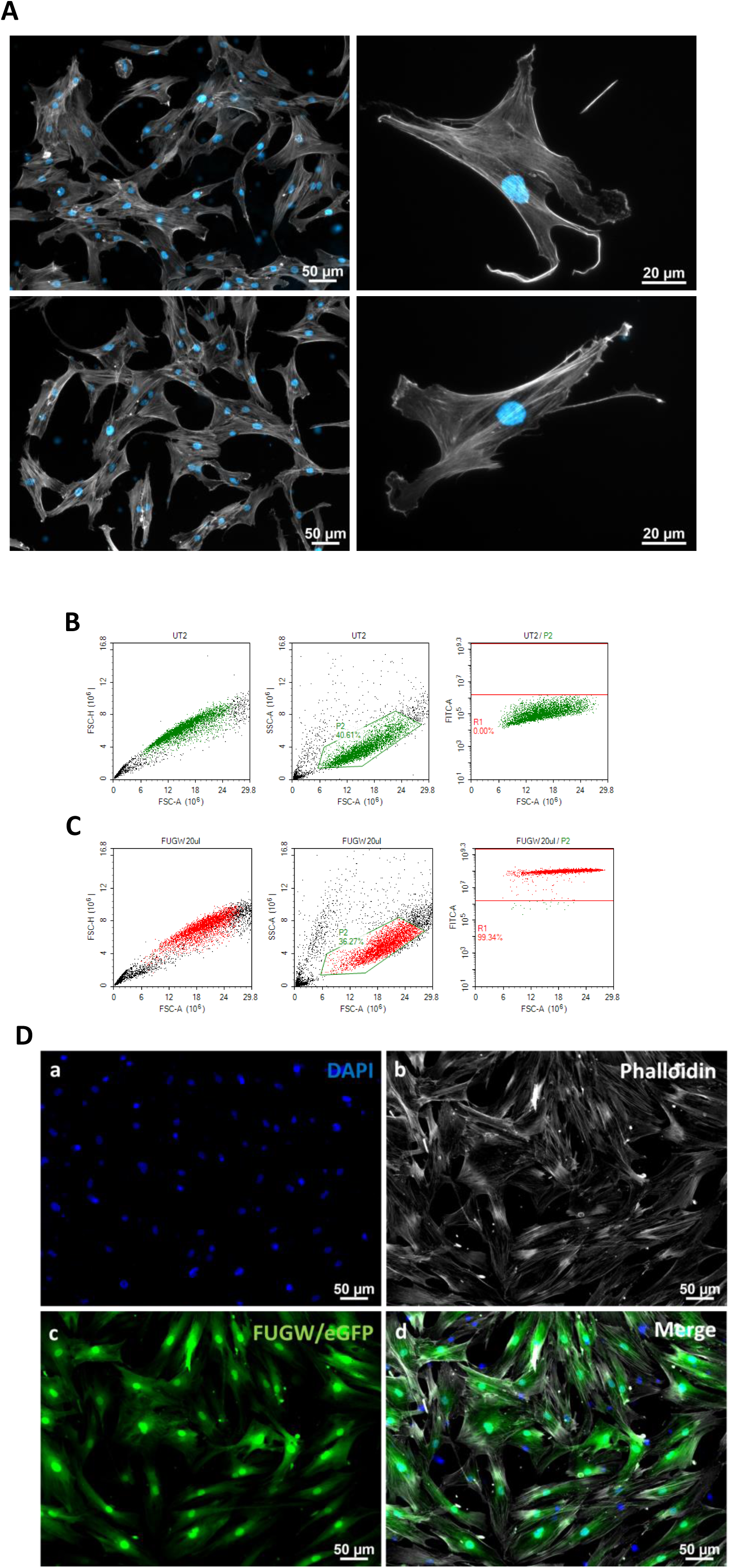
Representative images of passage 1 (P1) and 2 (P2) AT-MSC in-vitro cultures (A). Cells were visualized by epifluorescence microscopy following staining with DAPI and phalloidin (A). P2 AT-MSC cells were transduced with 20 μl of FUGW lentiviruses. After 3 days, cells were washed and trypsinized and thereafter, percentage of fluorescent cells was measured using FACS technology. Untreated AT-MSC cells did not display fluorescent expression (B top panels). The fraction of fluorescent AT-MSC cells following transduction with 20 μl of lentivirus FUGW 99.3% displayed by FITC-A detection within the gated population (C). Cell morphology and fluorescence expression were further analyzed using epifluorescence microscopy of transduced cells (D). Representative images of monolayer cultures of AT-MSC transduced with FUGW/GFP reporter PFA-fixed and stained with DAPI and phalloidin (D). Phalloidin staining displays the typical mesenchymal morphology of transduced cells (Db) and fluorescence expression is clearly visible (Dc). Scale bars: 50 µm (A-left panel, and D), and 20 µm (A-right panels).

**Supplementary Figure 2.**
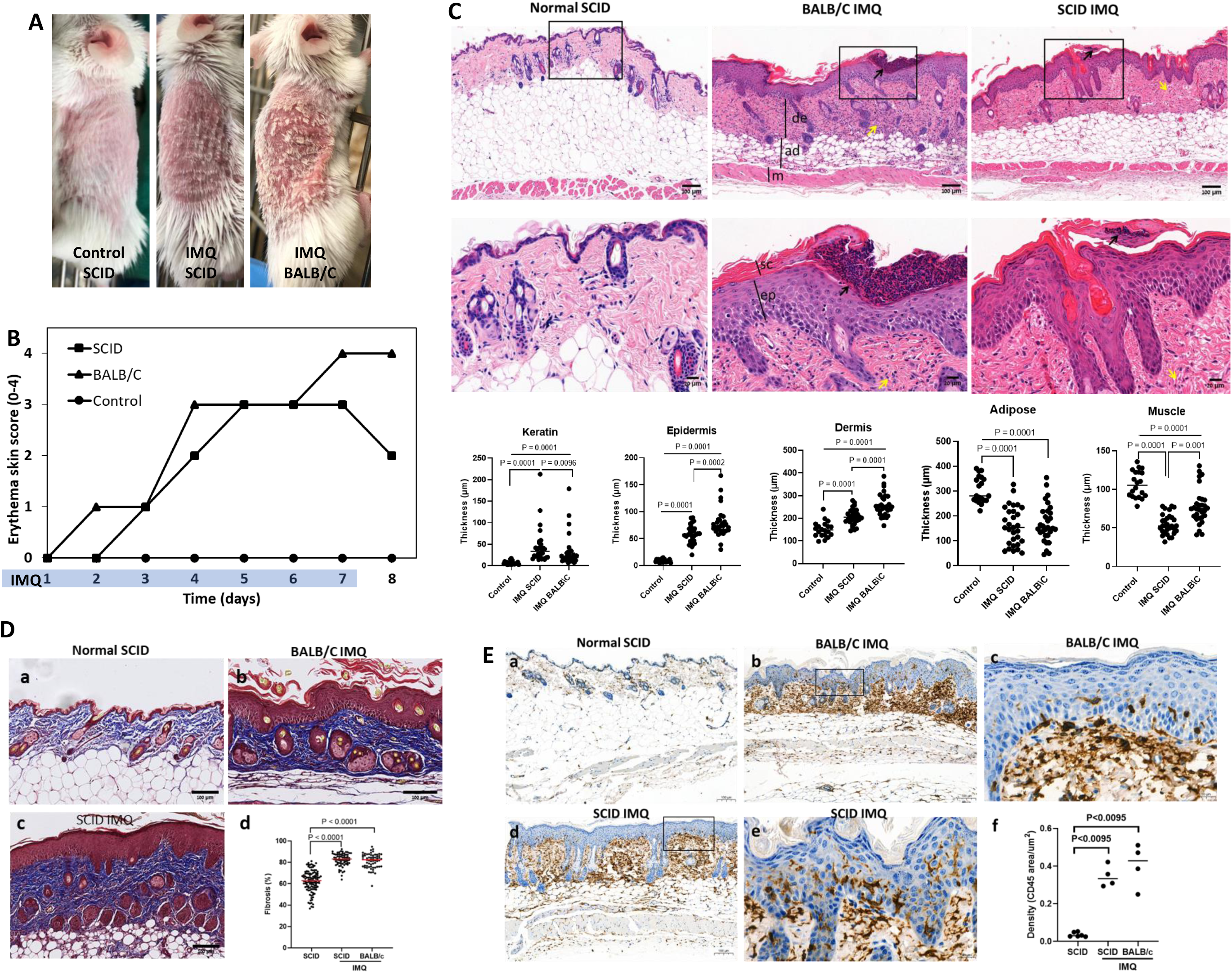
Induction of psoriatic-like skin disease in SCID mice. Female 8-12 weeks old BALB/c or SCID mice (A) were treated topically with imiquimod (IMQ) ointment once daily for 7 days and sacrificed on day 8 (B). Age and sex-matched SCID mice were used as normal control. FFPE skin tissues were sectioned and stained with H&E (C), Masson Trichrome (MT in D) and immunohistochemically (IHC) with ant-CD45 antibodies. Thickness of the stratum corneum (SC), epidermis (ep), dermis (de), adipose layer (ad), and muscle layer (m) was measured and quantified (C). Using MT staining dermal collagen was visualized and quantified (D). IHC staining using ant-CD45 demonstrated massively infiltration of the epidermis and dermis by immune cells in IMQ-treated mice (E). Images were acquired using a 3D HISTECH Pannoramic-250 microscope slide-scanner (3D HISTECH, Budapest, Hungary). Snapshots were taken with Case Viewer software (3D HISTECH, Budapest, Hungary). All quantitative analysis; thickness measurements (C bottom panels), collagen (Dd) and CD45 staining (Ef), were performed using ImageJ software package. Scale bars: 100µm (C top panels, D, and Ea,b&d), and 20µm (C bottom panels, and Ec&e).

**Supplementary Figure 3.**
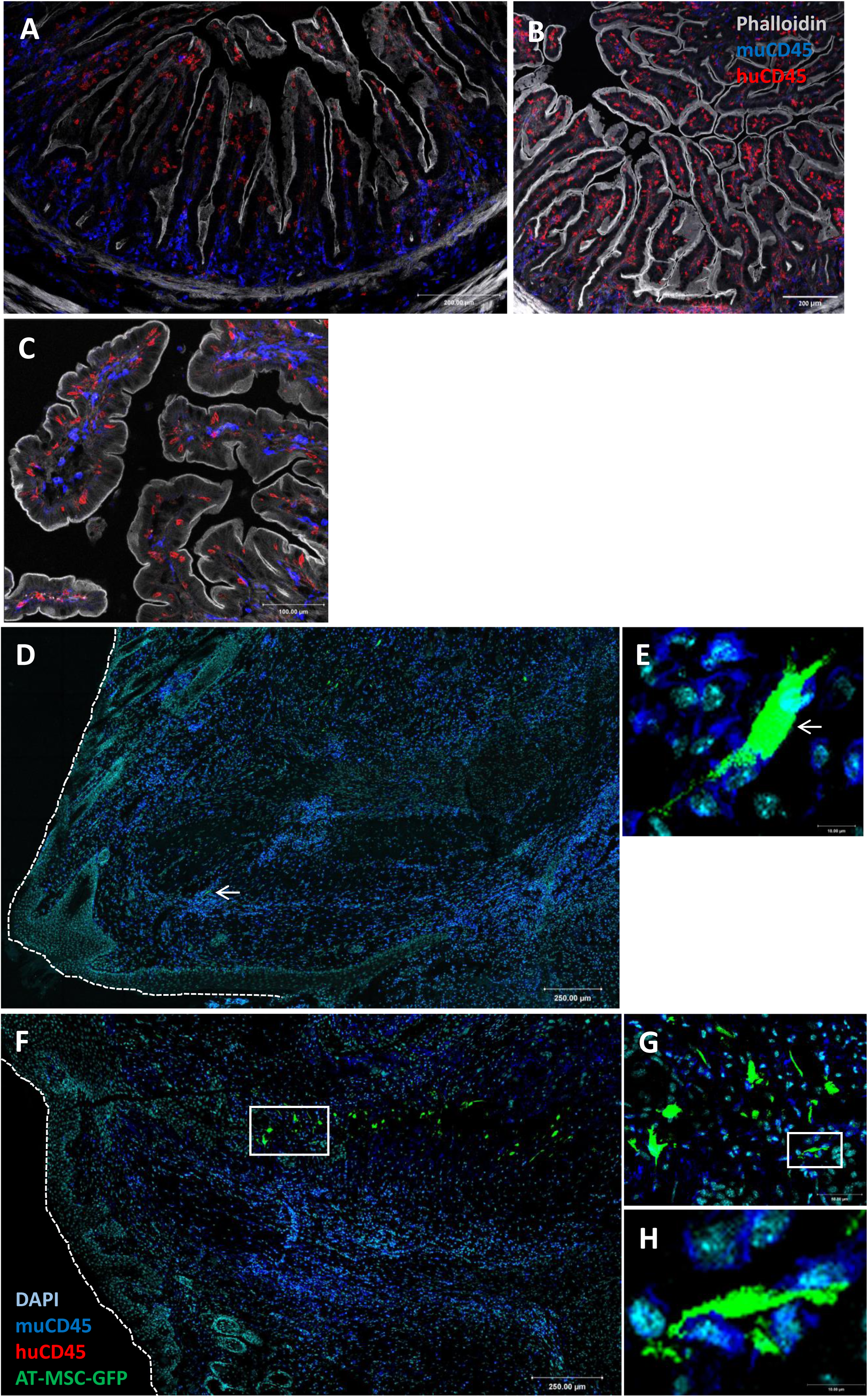
Human and murine CD45-positive immune cells populate non-treated steady state (A) and IMQ-LPS gut transplants (B-C). Fistula wound and fistula tract is populated by murine CD45-positive cells (D-H). Representative epifluorescence images of two edges of fistula wound and fistula tracts populated by reactive AT-MSC expressing eGFP (D and F), doted lines denote epidermal outer borders. huMSC expressing eGFP and murine CD45-positive immune cells (arrow in D and boxed area in F) were imaged in higher magnification (E and G-H). Scale bars; 200µm (A-B), 100µm (C), 250µm (D&F), 50µm (G) and 10µm (E &H).

**Supplementary Figure 4.**
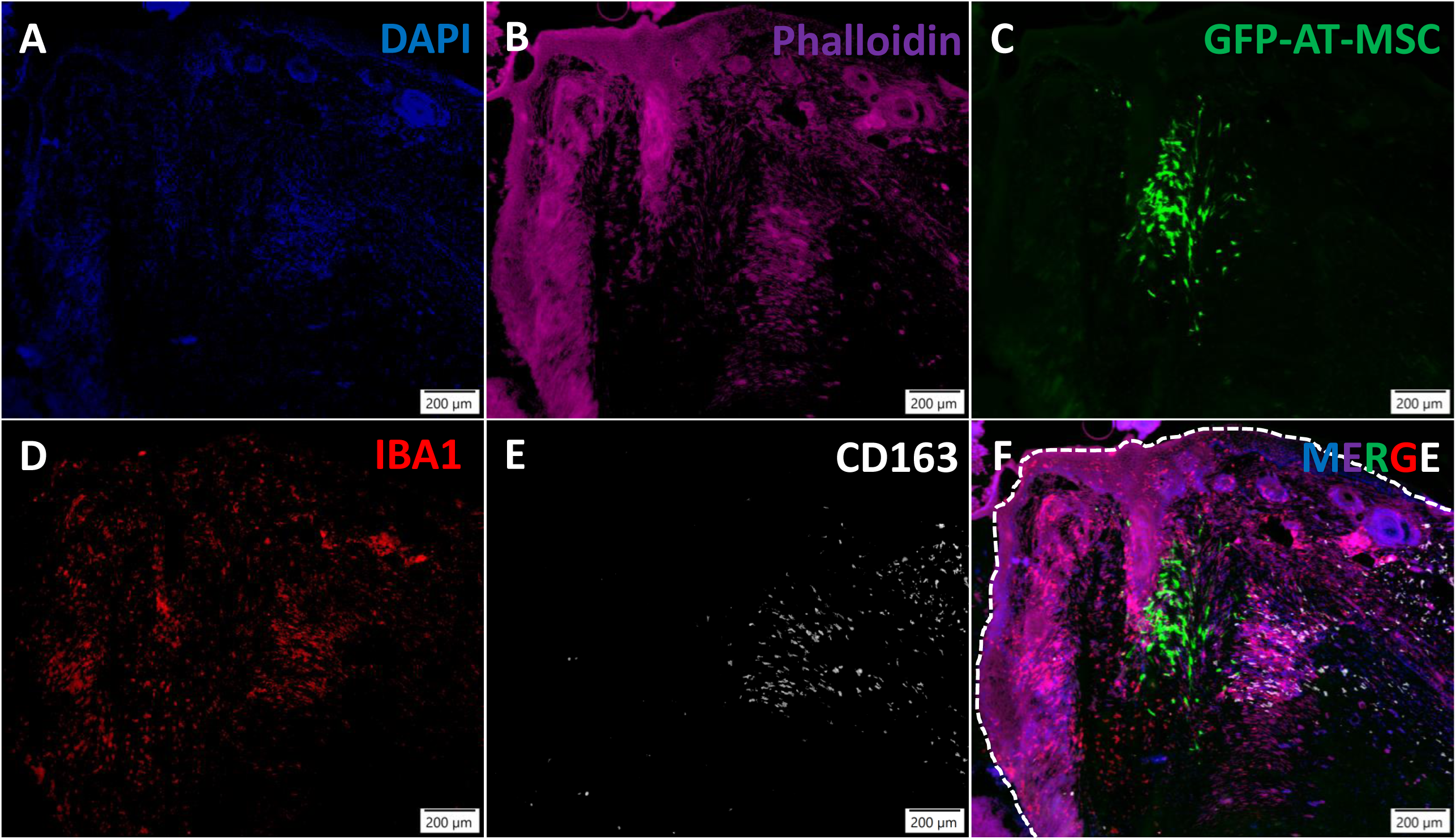
Individual fluorescence channels for Figure 6G. Representative image of AT-MSC treated fistula tract tissue stained with DAPI (blue in A and F), phalloidin (pink in B and F), anti-IBA1 (red in D and F) and anti-CD163 (white in E and F). Dotted line in F indicates outer border of epidermis. AT-MSC expressing eGFP are visible in C and F. Image was acquired using Olympus SLIDEVIEW VS200 scanner and snapshots were taken with OlyVIA v. 4.1.1 software. Scale bar; 200µm (A-G).

**Supplementary Figure 5.**
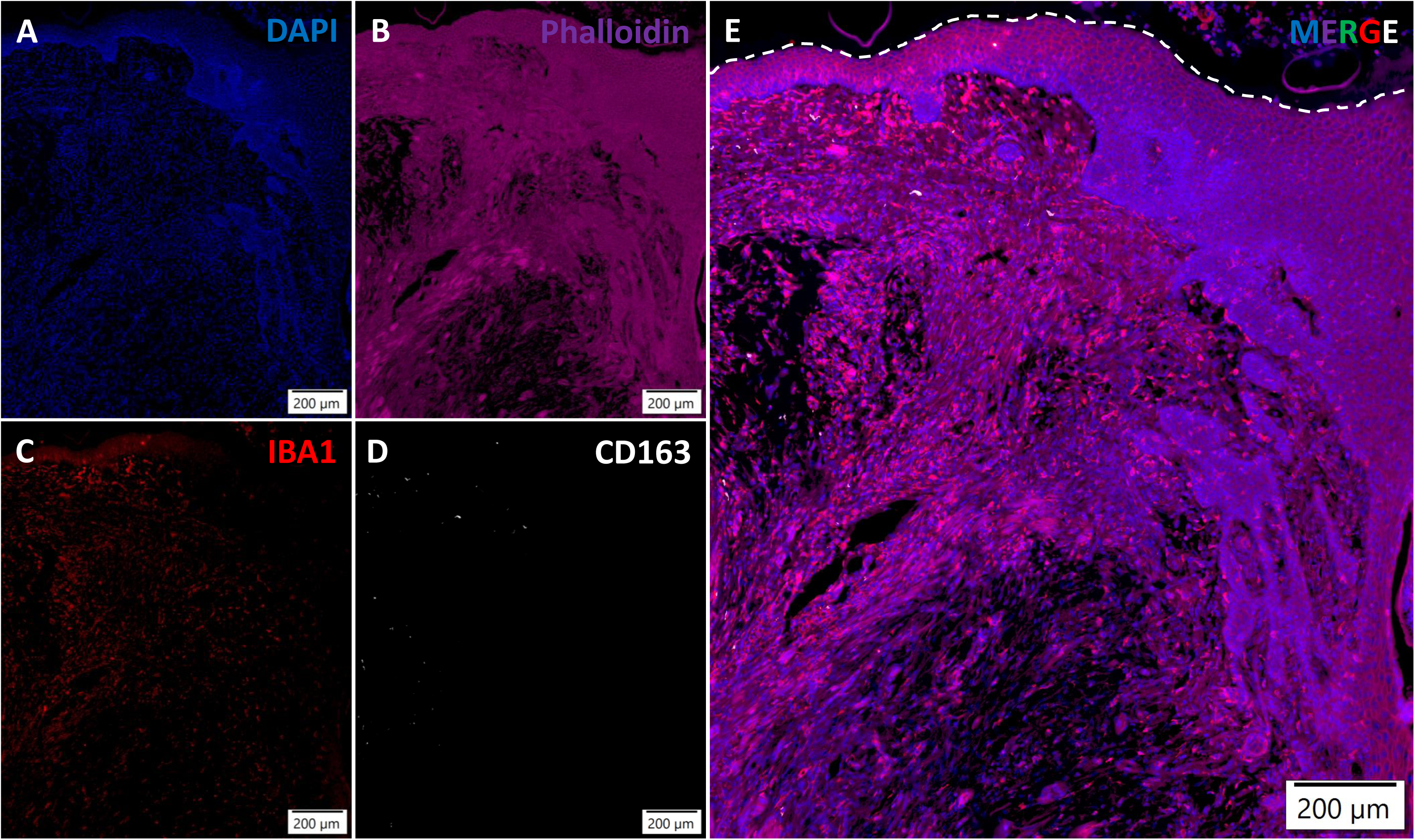
Individual fluorescence channels for Figure 6H. Representative image of non-treated control fistula tract tissue stained with DAPI (blue in A and F), phalloidin (pink in B and F), anti-IBA1 (red in D and F) and anti-CD163 (white in E and F). Dotted line in F indicates outer border of epidermis. Image was acquired using Olympus SLIDEVIEW VS200 scanner and snapshots were taken with OlyVIA v. 4.1.1 software. Scale bar; 200µm (A-E).

